# Cervicovaginal microbiome composition drives metabolic profiles in healthy pregnancy

**DOI:** 10.1101/840520

**Authors:** Andrew Oliver, Brandon LaMere, Claudia Weihe, Stephen Wandro, Karen L. Lindsay, Pathik D. Wadhwa, David A. Mills, David Pride, Oliver Fiehn, Trent Northen, Markus de Raad, Huiying Li, Jennifer B.H. Martiny, Susan Lynch, Katrine Whiteson

**Affiliations:** Dept. of Molecular Biology and Biochemistry, University of California, Irvine; Division of Gastroenterology, Dept. of Medicine, University of California, San Francisco, San Francisco; Dept. of Ecology and Evolutionary Biology, University of California, Irvine; Center for Microbiome Innovation, University of California, San Diego, San Diego; Dept. of Pediatrics and the Development, Health and Disease Research Program, College of Health Sciences, University of California, Irvine, California 92697; Dept. of Psychiatry and Human Behavior, College of Health Sciences, University of California, Irvine, California 92697; Dept of Food Science & Technology, University of California, Davis; Dept of Viticulture & Enology, University of California, Davis; Department of Pathology, University of California, San Diego, San Diego; Department of Medicine, University of California, San Diego, San Diego; West Coast Metabolomics Center, University of California, Davis; Environmental Genomics and Systems Biology Division, Lawrence Berkeley National Laboratory, Berkeley, California 94720; Department of Molecular and Medical Pharmacology, Crump Institute for Molecular Imaging, David Geffen School of Medicine, University of California, Los Angeles

## Abstract

**Background:** Microbes and their metabolic products influence early-life immune and microbiome development, yet remain understudied during pregnancy. Vaginal microbial communities are typically dominated by one or a few well adapted microbes, which are able to survive in a narrow pH range. In comparison to other human-associated microbes, vaginal microbes are adapted to live on host-derived carbon sources, likely sourced from glycogen and mucin present in the vaginal environment.

**Methods:** Using 16S rRNA and ITS amplicon sequencing, we characterized the cervicovaginal microbiomes of 18 healthy women throughout the three trimesters of pregnancy. Shotgun metagenomic sequencing permitted refinement of the taxonomy established by amplicon sequencing, and identification of functional genes. Additionally, we analyzed saliva and urine metabolomes using GC-TOF and LC-MS/MS lipidomics approaches for samples from mothers and their infants through the first year of life.

**Results:** Amplicon sequencing revealed most women had either a simple community with one highly abundant species of *Lactobacillus* or a more diverse community characterized by a high abundance of *Gardnerella*, as has also been previously described in several independent cohorts. Integrating GC-TOF and lipidomics data with amplicon sequencing, we found metabolites that distinctly associate with particular communities. For example, cervicovaginal microbial communities dominated by *Lactobacillus crispatus* also have high mannitol levels, which contradicts the basic characterization of *L. crispatus* as a homofermentative *Lactobacillus* species. It may be that fluctuations in which *Lactobacillus* dominate a particular vaginal microbiome are dictated by the availability of host sugars, such as fructose, which is the most likely substrate being converted to mannitol. Furthermore, indole-3-lactate (ILA) was also indicative of *L. crispatus* specifically. ILA has immunomodulatory properties through binding the human aryl hydrocarbon receptor (AhR), which may maintain the especially low diversity of *L. crispatus* dominated communities.

**Conclusions:** Overall, using a multi-‘omic approach, we begin to address the genetic and molecular means by which a particular vaginal microbiome becomes vulnerable to large changes in composition.

## Introduction

Vaginal microbes sustain important physiologies and produce metabolites that can directly or indirectly affect maternal health and infant development during pregnancy. Perturbations to early-life microbiomes and associated metabolic dysfunction have been linked with allergy and auto-immune diseases such as asthma [1–4]. For example, regular pre-natal and post-natal farm exposure, i.e., contact with a diversity of microbes during pregnancy and infancy, have been shown to reduce the incidence of chronic health diseases such as asthma and atopy [5]. Moreover, recent research has supported the idea of fetal programming, a term describing the process by which the maternal microbiota, as well as maternal antibodies, prepare the infant immune system for the post-natal onslaught of colonizing microbes [6]. Others have shown in mice that vaginal dysbiosis, induced by maternal stress, has the potential to negatively affect offspring metabolic profiles [7]. Thus maternal microbes, particularly those of the vaginal tract, are some of the first microbes the offspring will encounter and may be central to the study of early-life microbiome and immune development [8–12]. Indeed, a recent large-scale study of 2,582 women, over 600 of whom were pregnant (a subset of whom were longitudinally sampled), provided evidence for vaginal microbiome restructuring during pregnancy toward a *Lactobacillus-*dominated community [13]. This occurred early in gestation and was associated with a reduced vaginal microbiome metabolic capacity. Post-partum, irrespective of the mode of delivery, the vaginal microbiota resembled that of a gastrointestinal microbiome, likely due to microbial mixing during the birthing process [9], suggesting that both vaginal and gastrointestinal microbial seeding of the neonate occurs.

The human vaginal microbiome maintains low diversity in low pH conditions, and depends on host sugars as carbon sources, with less access to dietary and exogenous nutrients compared to the gut or the skin or the oral cavity. Historically, vaginal microbial communities have been stratified based on hierarchical clustering of the taxa composition [12]. Keystone species include *Lactobacillus crispatus* and *L. gasseri* which have been associated with maintenance of a simple vaginal microbiome by their production of bacteriostatic and bactericidal compounds (e.g. lactic acid and hydrogen peroxide) and maintenance of a low pH [14–16], numerically and functionally dominating their respective vaginal communities. A closely related species, *L. iners*, has been associated with health promoting benefits; however, its genome also encodes the capacity to promote microbiome perturbation by increasing vaginal pH and producing species specific virulence factors [14,17–19]. Bacterial vaginosis (BV), the most common gynecological condition in reproductive aged women [20], is characterized by the presence of a more diverse vaginal microbiome and associated with adverse pregnancy outcomes including preterm birth [21], endometritis [22, 23] and spontaneous abortion [24–27]. Recently, vaginal microbial transplants have been successfully implemented as a treatment for intractable BV [28]. Despite *L. crispatus* generally being regarded as a highly beneficial and dominant microbe throughout pregnancy, healthy women from different ethnic groups have markedly different species dominating the vaginal microbiome [15]. In fact, many healthy women that lack BV symptoms have vaginal microbiomes dominated by microbes that are associated with BV [29], suggesting that taxonomy alone is insufficient to predict health outcomes and that microbial activities, including metabolic productivity may offer a more contemporary view of microbiome function.

An untargeted, more global assessment of microbiomes and associated metabolites during pregnancy and early-life is lacking. To address this gap in knowledge, we collected saliva, urine, and cervical vaginal fluid (CVF) from 18 mothers during each trimester of pregnancy and saliva and urine from offspring through their first year of life. Specifically, we were interested in how maternal CVF microbiome profiles are associated with metabolomic assessments of the same samples. Furthermore, we had the opportunity to examine whether maternal saliva and urine metabolome profiles relate to those of the infant in the first year of life. Here we present DNA sequencing (amplicon and shotgun) and untargeted metabolomics to characterize microbial and metabolic features of the CVF microbiome throughout pregnancy to determine the composition of the vaginal microbiome from a cohort of healthy Caucasian and Hispanic women, longitudinally sampled throughout a healthy pregnancy.

## Methods

### Subject Information

Eighteen women were selected from a larger cohort recruited to address how maternal stress affects child development [30, 31] (Table 1). Inclusion criteria for the larger cohort included >18 years of age, singleton, intrauterine pregnancy, and non-diabetic. Additional inclusion criteria for this study were: normal pre-pregnancy body mass index, vaginal delivery, full-term pregnancy, breastfeeding, and no antibiotics for mother or baby. Generally, these eighteen women and their children represented healthy subjects with the most complete sample sets.

**Table 1:** Demographics of the eighteen mothers who participated in the study.

| **Maternal ID** | **Race-Ethnicity** | **Age** | **BMI (pre-pregnancy)** |
| --- | --- | --- | --- |
| 1018 | White Hispanic | 35 | 27.4 |
| 1062 | White Hispanic | 23 | 25.3 |
| 1088 | White non-Hispanic | 26 | 25.8 |
| 1089 | White non-Hispanic | 22 | 21.8 |
| 1103 | White non-Hispanic | 27 | 24.5 |
| 1111 | White Hispanic | 38 | 27.9 |
| 1120 | White non-Hispanic | 34 | 26.9 |
| 1126 | White Hispanic | 19 | 27.8 |
| 1137 | White Hispanic | 31 | 23.5 |
| 1146 | White non-Hispanic | 29 | 23.5 |
| 1151 | White non-Hispanic | 30 | 22.4 |
| 1157 | White non-Hispanic | 28 | 18.9 |
| 1180 | White non-Hispanic | 29 | 24.9 |
| 1191 | White Hispanic | 31 | 22.7 |
| 1198 | White non-Hispanic | 26 | 24.8 |
| 1201 | White non-Hispanic | 30 | 29.9 |
| 1202 | White Hispanic | 20 | 24.7 |
| 1222 | White non-Hispanic | 24 | 24.0 |

### Sample collection

Samples were collected at each trimester of pregnancy for women and through the first year of life for infants. At each timepoint, maternal saliva, urine and cervical vaginal fluid were collected. For infants, urine was collected at birth, six months, and twelve months whereas saliva was sampled at six months and twelve months of age. Maternal saliva was collected using a salivette collection kit, including a small cotton roll contained in a plastic container (Salimetrics, Carlsbad, CA). Mothers were instructed to place the cotton rolls in their mouth until saturated with saliva (approximately 1-3 minutes), and then reseal the swabs in plastic salivette tubes. Infant saliva was collected using Weck-Cel spears and a swab extraction tube system. Infants were allowed to suck on the spear for two minutes, ensuring saturation. Maternal urine was collected using a sterile collection cup. Infant urine was collected using an adhesive u-bag attached to the genital region of the infant. A minimum of 2 ml of urine was collected. Cervical vaginal fluid (CVF) was collected by placing three Dacron swabs into the cervix for ten seconds to achieve saturation. Each swab was then placed in a plastic vial with 500 μl of sterile PBS. All samples were initially stored at −20°C and then saliva, CVF, and infant urine were subsequently moved to −80°C storage.

### Metabolomics

Prior to processing, samples were thawed from −80°C storage. Fifty microliters of each sample were subjected to gas chromatography time-of-flight mass spectrometry (GC-TOF) [32] and liquid chromatography accurate mass mass-spectrometry (LC-MS/MS, lipidomics). Urine, saliva, and CVF from each time point were sent to the West Coast Metabolomics Center (WCMC) for untargeted metabolomics. GC-TOF metabolites were extracted with a mixture of 3:3:2 acetonitrile:isopropyl alcohol:water according to standard operating procedures from the Fiehn Lab at the WCMC [33]. LC-MS/MS samples were extracted using a variant of the Matyash method [34]. Data were acquired for complex lipids in positive and negative electrospray mode on a Waters CSH column and an Agilent 6530 QTOF mass spectrometer [34]. Metabolites were identified by retention time MS/MS matching using MassBank of North America (http://massbank.us) and NIST17 libraries.

### High-throughput metabolomics

All urine, saliva and CVF were analyzed using Matrix Assisted Laser Desorption Ionization (MALDI) Imaging Mass Spectrometry (MSI) for high-throughput untargeted metabolomics. Extracted samples in 3:3:2 acetonitrile:isopropyl alcohol:water were diluted 1:2 in water in 384 well plates. Next, an equal volume of MALDI matrix (20 mg/mL of 1:1 2,5-Dihydroxybenzoic acid and α-cyano-4-hydroxycinnamic acid in 1:3 (v/v) H2O:MeOH+0.2% formic acid) was added. Samples were printed onto a stainless steel blank MALDI plate using an ATS-100 acoustic transfer system (BioSera) with a sample deposition volume of 10nl. Samples were printed in clusters of four replicates, with the microarray spot pitch (center-to-center distance) set at 900 um). MS-based imaging was performed using an ABI/Sciex 5800 MALDI TOF/TOF mass spectrometer with laser intensity of 3,500 over a mass range of 50–3,000 Da. Each position accumulated 20 laser shots. The instrument was controlled using the MALDI-MSI 4800 Imaging Tool. Surface rasterization was oversampled using a 75 um step size. Average ion intensity for all reported ions was determined using the OpenMSI Arrayed Analysis Toolkit (OMAAT) software package [35].

### DNA extraction

Cervical brushes were resuspended in PBS. Aliquots of 100-200 μl were added to lysing matrix E tubes pre-aliquoted with 500 of hexadecyltrimethylammonium bromide (CTAB) DNA extraction buffer and incubated at 65°C for 15 minutes. An equal volume of phenol:chloroform:isoamyl alcohol (25:24:1) was added to each tube and samples were homogenized in a Fast Prep-24 homogenizer at 5.5 m/s for 30 seconds. Tubes were centrifuged for 5 minutes at 16,000 x *g* and the aqueous phase was transferred to individual wells of a 2 ml 96-well plate. An additional 500 µl of CTAB buffer was added to the lysing matrix E tubes, the previous steps were repeated, and the aqueous phases from paired extractions were combined. An equal volume of chloroform was mixed with each sample, followed by centrifugation at 3000 x *g* for 10 minutes. The aqueous phase (600 µl) was transferred to a clean 2 ml 96-well plate, combined with 2 volume-equivalents of polyethylene glycol (PEG) and stored overnight at 4°C to precipitate DNA. Plates were centrifuged for 60 min at 3000 x *g*. DNA pellets were washed twice with 300 µl of 70% ethanol, air-dried for 10 minutes and re-suspended in 100 µl of sterile water. DNA was quantified using the Qubit dsDNA HS Assay Kit and diluted to 10 ng/µl, when possible. Although DNA was extracted from CVF, attempts to extract DNA from saliva were unsuccessful, potentially due to the storage swabs trapping the biomaterial.

### Amplicon gene sequencing

To amplify the V4 region of the bacterial 16S rRNA gene, 10 ng of DNA template was combined with PCR master mix (0.2 mM dNTP mix, 0.56 mg/ml BSA, 0.4 uM Illumina adapter sequence-tagged forward primer (515F) [36], 0.025 U/μl Taq DNA polymerase) and 0.4 uM barcode-tagged reverse primers (806R) then amplified in triplicate 25 μl reactions, along with a no-template control, for 30 cycles (98°C for 2 min; 98°C for 20 sec, 50°C for 30 sec, 72°C for 45 sec; repeat steps 2-4 29 times; 72°C for 10 min). PCR conditions were identical for ITS2 amplification (primer pair fITS7 (5′-GTGARTCATCGAATCTTTG-3′) and ITS4 (5′-TCCTCCGCTTATTGATATGC-3′)) except for the annealing temperature, which was 52°C. Triplicate reactions were combined and purified using the SequalPrep Normalization Plate Kit (Invitrogen) according to manufacturer’s specifications. Purified amplicons were quantified using the Qubit dsDNA HS Assay Kit and pooled at equimolar concentrations. The amplicon library was concentrated using the Agencourt AMPure XP system (Beckman-Coulter), quantified using the KAPA Library Quantification Kit (KAPA Biosystems) and diluted to 2nM. Equimolar PhiX was added at 40% final volume to the amplicon library; the 16S rRNA amplicon pool was sequenced on the Illumina NextSeq 500 Platform on a 153bp x 153bp sequencing run, and the ITS2 amplicon pool was sequenced on the Illumina MiSeq platform on a 290bp x 290bp run.

### Shotgun metagenomics sequencing

Sequencing libraries were prepared using the Illumina Nextera kit and methods described in Baym et al. [37]. Briefly, DNA from each sample was diluted to 0.5ng/µl and tagmented with the Nextera enzyme (Illumina) for 10 min at 55°C. Following tagmentation, each sample received 1 µl forward and 1 µl reverse barcodes, which were added via PCR using Phusion DNA polymerase (New England BioLabs). After PCR, the libraries were cleaned of smaller DNA fragments, using AMPure XP magnetic beads (Beckman-Coulter), and pooled by concentration. Libraries were quantified using the Quanti-iT PicoGreen dsDNA kit (Thermo Fischer Scientific), and DNA was run on a gel to check fragment size. These libraries were loaded onto the Illumina Next-Seq 500 at 1.8 picomolar concentrations and Illumina’s mid-output kit for 75 bp paired-end sequencing.

### OTU Table Generation

Raw sequence data were converted from bcl to fastq format using bcl2fastq v2.16.0.10. Paired sequencing reads with a minimum overlap of 25bp were merged using FLASH v1.2.11. Index sequences were extracted from successfully merged reads and demultiplexed in the absence of quality filtering in QIIME (Quantitative Insights Into Microbial Ecology, v1.9.1), and reads with more than two expected errors were removed using USEARCH’s fastq filter (v7.0.1001). Remaining reads were de-replicated, clustered into operational taxonomic units (OTUs) at 97% sequence identity, filtered to remove chimeric sequences, and mapped back to OTUs using USEARCH v8.0.1623. Taxonomy was assigned with the most current Greengenes database for bacteria [36] (May 2013), and UNITE vers. 6 for fungi [38]. OTUs detected in Negative Extraction Controls (NECs) were considered potential contaminants and filtered by subtracting the maximum NEC read count from all samples; any remaining OTU with a total read count less than 0.001% of the total read count across all samples was removed. Sequencing reads were rarefied to an even depth (65,029 reads for 16S; 91,232 reads for ITS2). To maximize similarity between the raw and rarefied OTU tables, random subsampling was performed at predefined depths 100 times, and the most representative subsampled community was selected based on the minimum Euclidean distance to the other OTU vectors generated for each sample.

### 16S rRNA Gene Analysis

Alpha-diversity indices and Bray-Curtis dissimilarity matrices were generated in QIIME [39]. Linear outcomes were assessed by linear mixed-effects (LME) modeling to adjust for repeated measures using the nlme package [40] in the R environment [41]. Variables of p<0.05 were considered statistically significant. Data was visualized using Tableau and Adobe Illustrator unless otherwise noted.

### Metagenomic Analysis

Raw sequences were filtered using Prinseq v0.20.4 [42] to filter out sequences that had a mean quality score of 30 or less. Human DNA was next filtered out by aligning the filtered reads to the human genome (hg38) using Bowtie2 v2.2.7 [43], and keeping the reads that failed to align (mean 208,225 reads or 10.24% of quality filtered reads per sample). To analyze functional potential, the reads were run through HUMAnN2 v0.1.9 [44] using default parameters and differences in pathway abundances were analyzed using LEfSe [45]. These reads were also cross assembled using SPAdes v3.8.2 [46]. Each sample was then mapped to this cross-assembly using Bowtie2, and samples from the same subject were merged together using Samtools v1.9 and the resulting bam files and the cross-assembly were imported into Anvio4 [47]. Taxonomy was assigned to each gene call using Kaiju [48] which subsequently informed a more accurate metagenomics binning of the most abundant microbes present.

### Statistical Analysis

Unless otherwise noted, statistics were done using the ecological statistics program Primer-e v7 [49]. Metabolic data were normalized in Primer-e by dividing by sum total for each sample. The specific programs used in Primer-e were permutational multivariate analysis of variance (PERMANOVA) and distance-based linear models (DistLM), the former of which calculates the significance and variance explained by a given factor and the latter determining which environmental variables correlate with the biological (microbiome) data. RFPermute [50], an R package for permutated random forests, was also performed to determine which annotated GC metabolites were indicative of microbial composition. PERMANOVA also partitions variance based on each factor, which is done in Primer-e by dividing the factor estimate by the sum total estimates of components of variation (ECoV). Traditional R^2^ values were also calculated by dividing the sum of squares by total sum of squares. LMEs were carried out as described above; R^2^ values for linear mixed models was calculated using the MuMIn package in R [51]. Relate tests (analogous to Mantel tests), were used to compare GCMS and LCMS data. Bray Curtis distances were used for all distance-based analyses. To consider repeated measures, linear mixed effects modeling (nlme package in R) was used to analyze stability of the microbiome and metabolome through time.

## Results

### Description of cohort and data obtained from samples

Saliva, urine, and cervical vaginal fluid (CVF) were collected from eighteen women, at early, middle and late pregnancy with the gestational age range of the included women at each timepoint (Fig. 1). At the time of enrollment into the cohort, the average woman’s age was 27.8 years old and average pre-pregnancy BMI was 24.8 (Table 1). The cohort was 39% white Hispanic and 61% non-Hispanic white; there were no significant differences in BMI (T-test, P=0.28) or age (T-test, P=0.89) between ethnic groups. Saliva and urine were collected at indicated intervals from each infant up until one year of age (Fig. 1). Saliva, urine, and CVF samples were subjected to metabolomics analysis whereas only CVF was used for sequence analysis. Sequence analysis included amplicon-based sequencing of the 16S rRNA gene (bacteria & archaea) and ITS2 (fungi) loci, and shotgun metagenomic sequencing of the entire microbial community.

**Figure 1:**
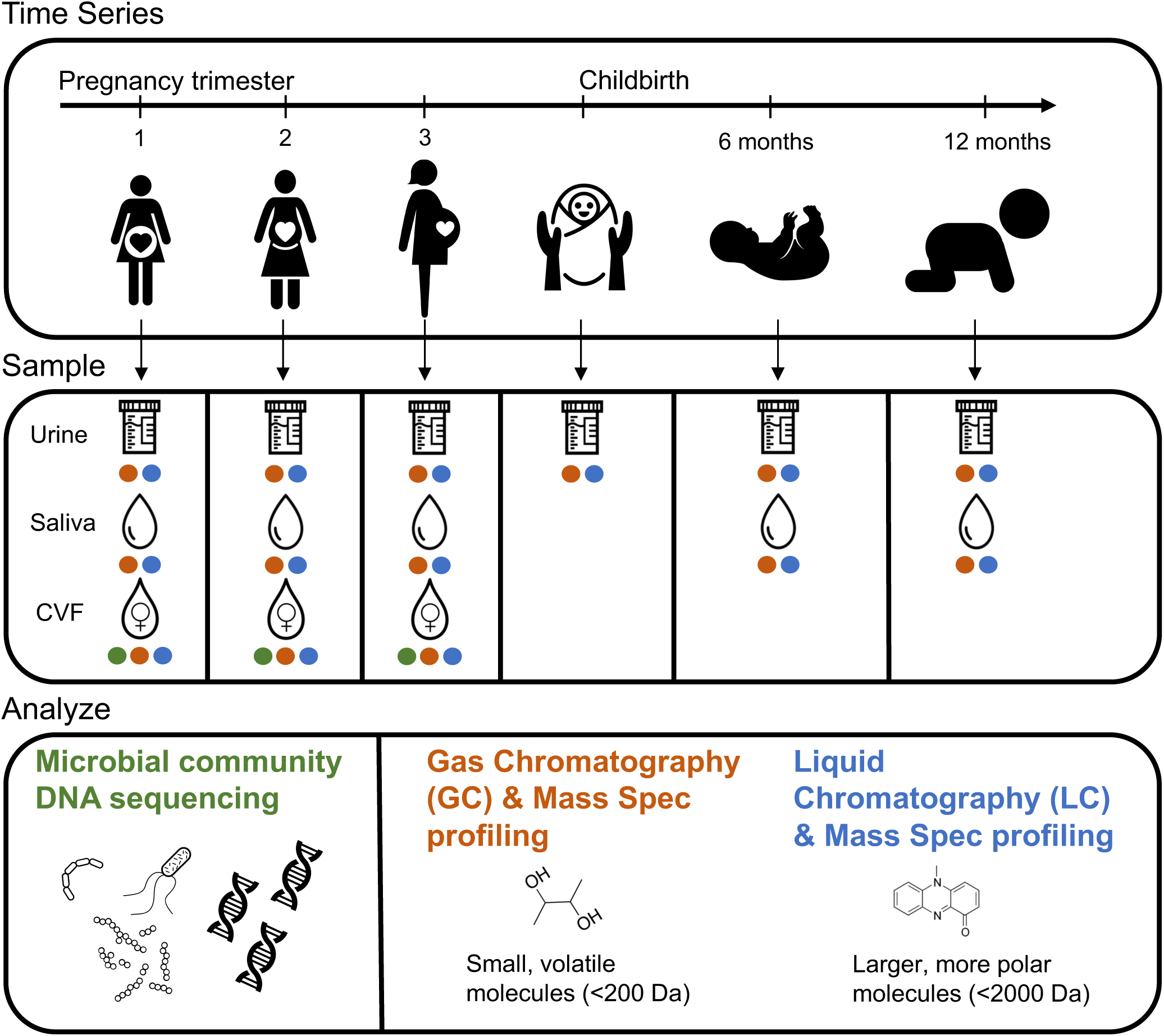
Study outline. Eighteen women were sampled throughout pregnancy and their offspring were sampled at birth, six, and 12 months of age. Samples collected were urine, saliva, and cervical vaginal fluid (CVF) for the mothers, and urine and saliva for the children. CVF was sequenced using shotgun metagenomics and amplicon sequencing. All samples were analyzed using GC-TOF and lipidomics.

### Vaginal microbiota support high abundances of Lactobacillus and Bifidobacteriaceae throughout pregnancy

16S rRNA gene sequencing stratified cervical samples into those where the most abundant taxon was *Lactobacillaceae* (34/42, 81%) or *Bifidobacteriaceae* (8/42, 19%) (Fig. 2A). The bacterial taxa in samples with abundant *Bifidobacteriaceae* were significantly more evenly distributed (LME, P=0.001, Fig. 2B), with *Gardnerella vaginalis* being the most abundant in seven of eight samples (88% relative abundance) and a *Shuttleworthia* taxon being most abundant in one sample (at 23% relative abundance). In samples where *Lactobacillaceae* had the highest abundance, a single taxon comprised 50% or more of the sequencing reads (27/34, 79%). *L. iners* was the most abundant taxon detected in 14 samples from 7 subjects, with a median relative-abundance of 79%. In twelve samples from 7 subjects, a *Lactobacillus* taxon, putatively identified as *L. crispatus* through metagenomic sequencing (Supp. Fig. 1), had a median relative abundance of 96%, and persisted at a relative abundance greater than 90% in subjects 1088, 1120, and 1191. Altogether, the most abundant 3 taxa (*L. iners* (OTU_1), *L. crispatus* (OTU_2), and *G. vaginalis* (OTU_3) comprised 66% of the total bacterial sequencing reads.

**Figure 2:**
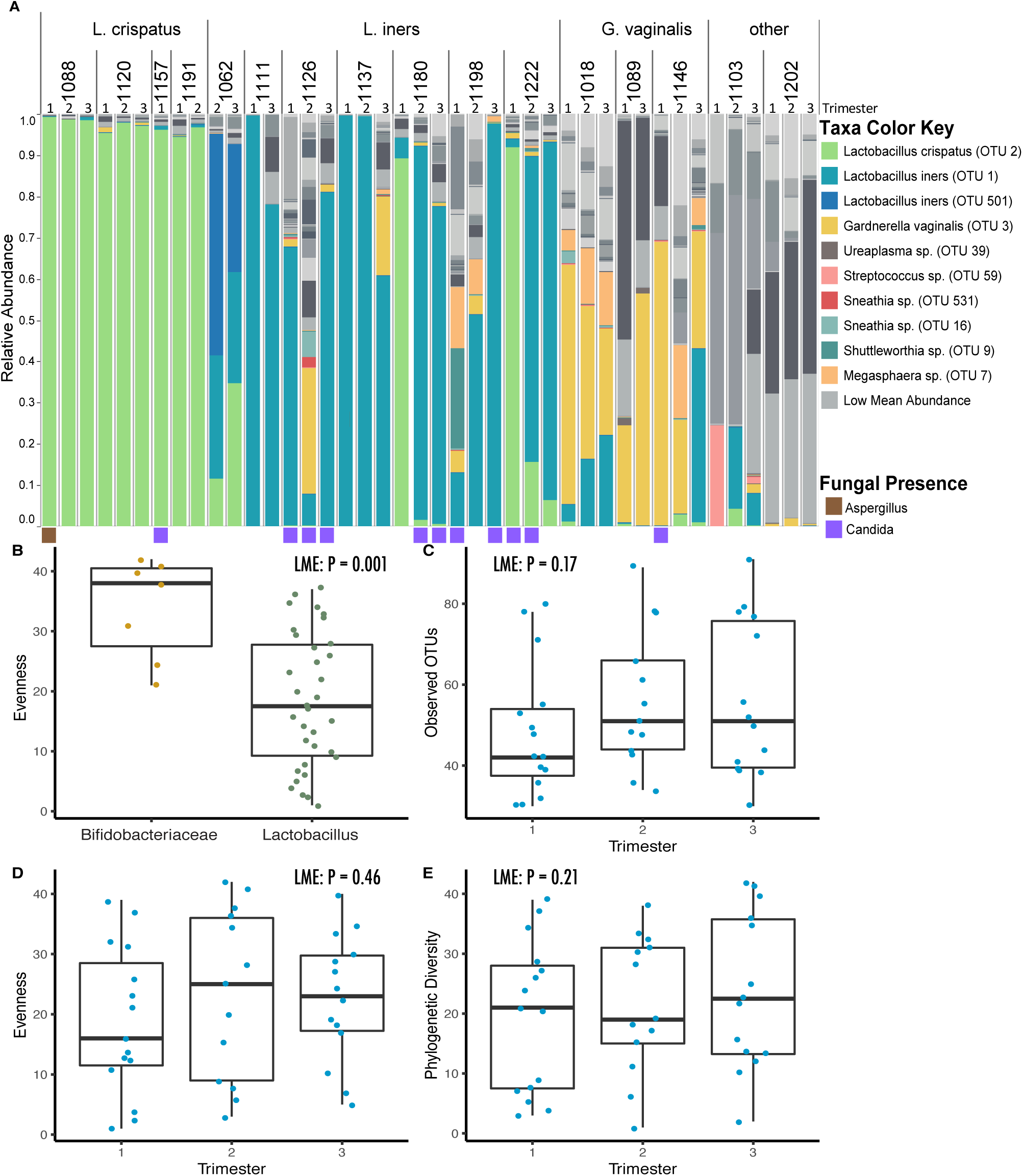
Taxonomy and alpha diversity of vaginal microbiomes during pregnancy. A) Relative abundance plot of operational taxonomic units, from 16S amplicon data, grouped together by individual. Each individual is clustered into a larger category defined by the dominating microbe. Linear mixed-effects models (LME) were done on the alpha diversity metrics to account for repeated measures in the data. B) Evenness between samples dominated by *Lactobacilli* is significantly lower than samples dominated by *Bifidobacteriaceae.* C) No significant change in the observed OTUs between the trimesters of pregnancy and likewise D) there was not change in evenness or E) phylogenetic diversity throughout pregnancy.

Whole genome shotgun sequencing produced, on average, 2.4 million reads per sample, which decreased to an average of 215,118 reads per sample following removal of reads that aligned to the human genome. Thirteen individuals produced sufficient sequence reads for taxonomic assignment, which was concordant with 16S rRNA gene sequence results (Supp. Fig. 1). Most of the reads classified as *L. crispatus* or *L. iners* mapped to a single metagenomic assembled genome, with completeness of 95.7% and 97.1% and redundancy of 0% and 1.4% respectively.

Twelve samples from seven subjects produced ITS2 sequences (Fig. 2a); we do not have quantitative data characterizing the abundance of bacterial or fungal biomass. Eleven samples from six subjects contained species of *Candida*, classified as *C. albicans* (Supp. Fig. 2), the most abundant fungal taxon in these data. Subject 1088 was the only participant to deviate from this trend, with a high relative abundance of *Aspergillus* during the first trimester of pregnancy (Supp. Fig. 2).

Alpha diversity indices based on 16S rRNA, and ITS2 data when available were compared across trimesters. While some subjects exhibited qualitative evidence of compositional shifts in vaginal microbiota with advancing gestation (Fig. 2A), we did not observe a significant difference in bacterial richness (number of observed OTUs) (LME, p = 0.17), evenness (Pielou’s evenness index) [52] (LME, p = 0.46) or phylogenetic diversity (LME, P=0.21)] across trimesters (Fig. 2C-E).

### Highly abundant bacterial taxa were significantly associated with community composition

An nMDS plot of Bray-Curtis dissimilarities showed that vaginal communities clustered by their most abundant bacterium (Fig. 3A). This association between the most abundant bacterial taxa and microbial composition of the sample was significant and explained more than half of the variance using PERMANOVA (R^2^= 56%, p= 0.0001). To account for repeated measures from longitudinal samples from the same individual we also performed an LME, which required dimensional reduction (LME, R^2^=69%, p < 0.0001). Communities with abundant *L. crispatus* were more similar to each other, sharing more than 90% similarity, in comparison to communities where a different bacterial species was most abundant. While some individuals exhibited a relatively stable microbial community over time, others (6/18) experienced shifts in composition, resulting in a statistically significant change in Bray-Curtis dissimilarity on PC axis-1 between trimesters (Fig. 3B).

**Figure 3:**
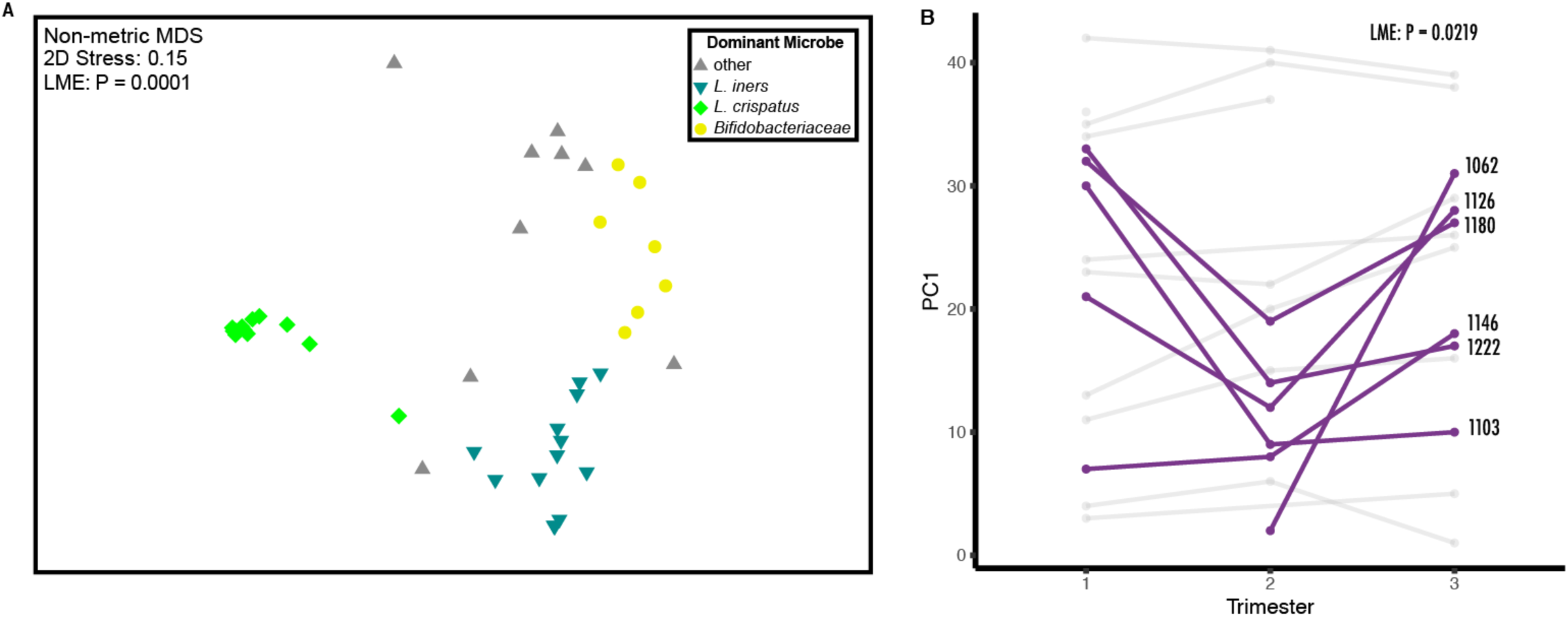
Ordination of vaginal microbiomes during pregnancy. A) Non-metric multidimensional scaling (nMDS) of Bray Curtis dissimilarity between vaginal microbiomes of mothers. Color indicates the most abundant microbe within the microbial community. The most abundant microbe in the community plays a statistically significant role in the composition of the community (LME, R^2^=69%, p < 0.0001) B) Some participants experienced large, significant shifts in their microbiomes throughout the trimesters of pregnancy.

The six subjects with *Candida* detected in at least one of two longitudinally-paired samples displayed a significant increase in inter-sample Bray-Curtis dissimilarity in their bacterial profiles over that interval (e.g. the intervals between trimester 1-2, 2-3, or 1-3), suggesting the presence of *Candida* may be associated with greater shifts in bacterial composition than those who had no *Candida* detected (Supp. Fig. 2B).

### Metabolites have strong associations with vaginal microbial community structures

Using GC-TOF MS, we detected 330 metabolites from urine, saliva, and CVF with 133 identified compounds. In the same samples, 1946 metabolites were also detected by LC-QTOF MS/MS (lipidomics, Supp. Table 1), with an additional 353 identified compounds. The CVF metabolome as assessed by both mass spectrometry methods did not significantly differ across trimesters (LME, GC-TOF MS: p = 0.6378, LC-MS/MS: p = 0.3942). This stability was even true for the subset of individuals who exhibited shifts in microbiota composition over trimesters (LME, GC-TOF MS: p = 0.6594, LC-MS/MS: p = 0.2482). CVF samples dominated by distinct bacteria exhibited significant differences in metabolic profiles (PERMANOVA, R^2^= 12%, p= 0.0195). A constrained, distance-based ordination plot recapitulated 67% of the community variation observed in the vaginal microbiota (Fig. 4). Superimposed on the ordination plot are GC-TOF predictor metabolites, calculated using the DistLM program in Primer-e. Indole-3-lactate (ILA) accounted for 27% of variation observed in the vaginal microbiota data, and was found to be more abundant in vaginal microbiota with abundant *L. crispatus* (Fig. 4B). Mannitol was also more abundant in samples dominated *L. crispatus* (Fig. 4C). In parallel we found that a pathway for mannitol production is also more abundant in shotgun metagenomic datasets of CVF samples dominated by *L. crispatus* (Supp. Fig. 3). This linear model identified the top ten annotated GC-TOF metabolites that were associated with variation in the microbial community composition are shown in Figure 4a; these ten metabolites together might explain almost 57% of the total variation in the microbial community composition. A permutated random forest recapitulated what we found in the DistLM, identifying mannitol and indole-3-lactate as two of the top variables of importance, specifically for distinguishing microbiomes with high abundances of *L. crispatus* (Supp. Fig. 4). To explore the ability to analyze the metabolome in high-throughput, the same sample sets were analyzed using Matrix Assisted Laser Desorption Ionization (MALDI) Imaging Mass Spectrometry (MSI) (Supp. Table 1). Detected ions by MALDI were compared to those identified by GC-MS and LC-MS, and found that ∼55% of the metabolites identified had corresponding ions in the MALDI analysis (Supp. Table 1).

**Figure 4:**
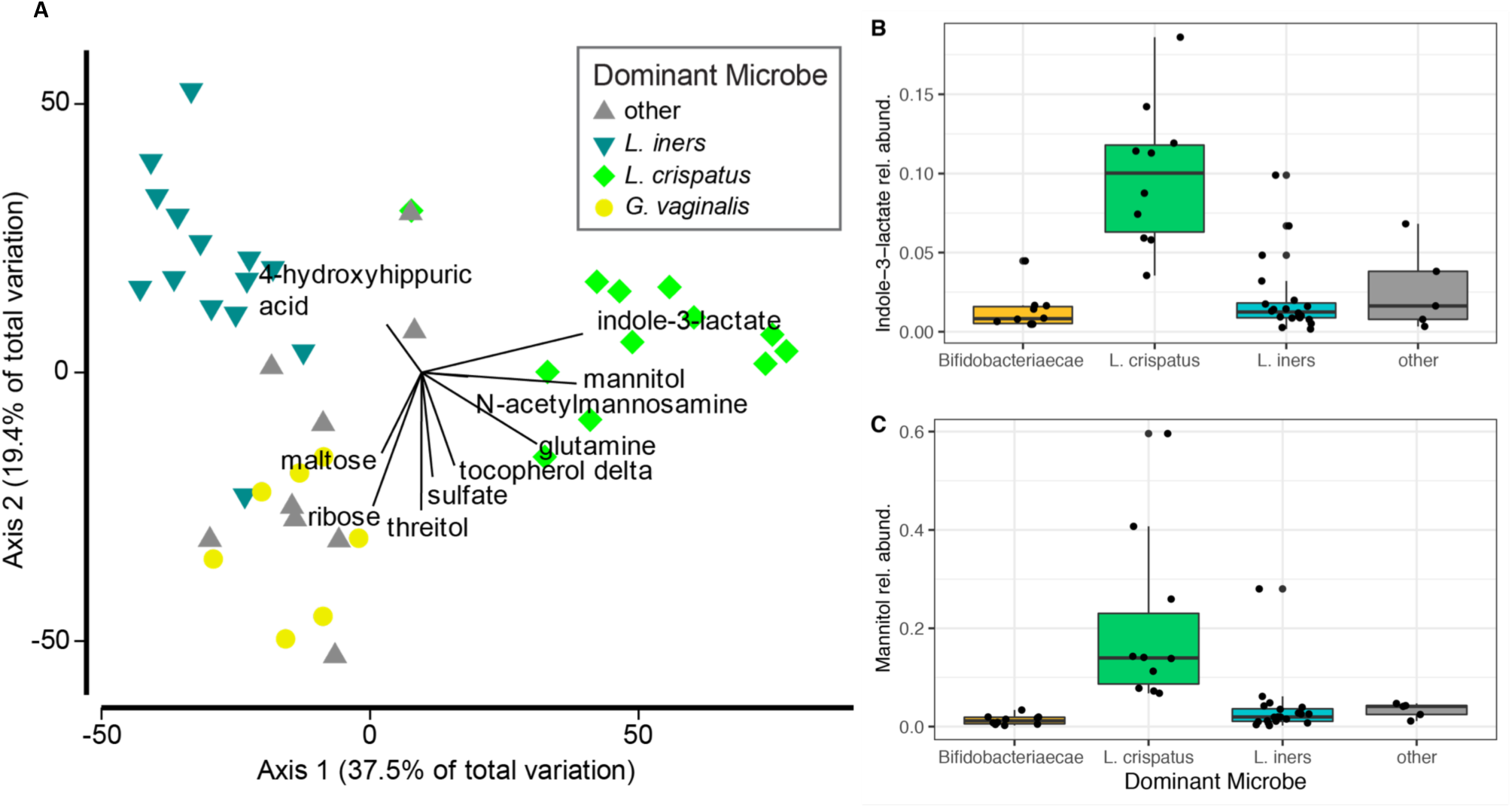
Relationship between vaginal microbes and metabolites. A) Distance-based linear model recapitulates the relationship between the vaginal microbiomes of these subjects. Superimposed are vectors showing which annotated GC-TOF molecules are best correlated with these microbial communities. Length and direction of vectors correspond to the strength of the association between the metabolite and the microbial communities. Boxplots show the raw abundance of B) indole-3-lactate and C) mannitol.

### Metagenomics and functional potential of communities

Distinct functions were associated with each of the vaginal microbial community clusters (PERMANOVA, R^2^= 70%, p= 0.0001, Fig. 5). LEfSe identified several pathways that differed between *L. crispatus* and *G. vaginalis*, in particular, an enrichment of ammonia assimilation genes in *G. vaginalis* (Supp. Fig. 3). Genes involved in mannitol metabolism were enriched in communities where *L. crispatus* was highly abundant (Supp. Fig. 5). Searching the PATRIC database of all sequenced *L. crispatus* (64 genomes), *L. iners* (22 genomes), and *G. vaginalis* (127 genomes) strains revealed annotated genes for mannitol usage and transport for *L. crispatus*, but not for *L. iners* or *G. vaginalis*.

**Figure 5:**
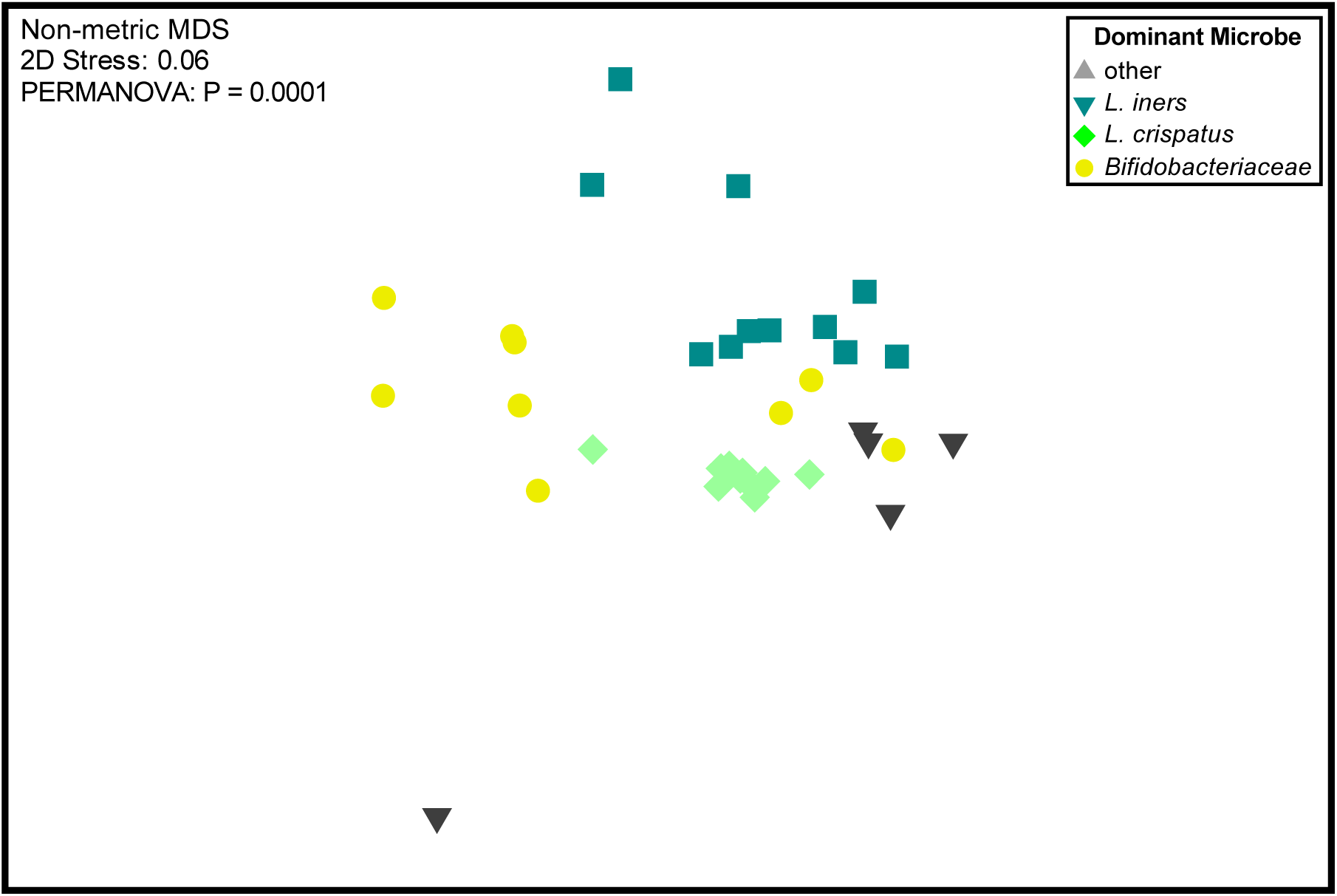
Ordination of functional pathways within the vaginal microbiome. An nMDS of HUMAnN2 analysis, examining the abundance of pathways in each microbiome. Vaginal microbiomes have functions that are indicative of the most abundant microbe present in the samples.

### Mothers and infants have significantly different saliva and urine metabolomes

Maternal and infant saliva and urine metabolomes were assessed with both GC-TOF and LC-MS/MS (lipidomics) in order to study the relationship between maternal and infant metabolomic compositions during early life (see Fig. 1). PERMANOVA analysis of lipidomics data from saliva samples showed the largest difference between mothers and offspring (PERMANOVA, R^2^= 69%, p = 0.001, Supp. Fig. 6A). A subset of 50 lipidomics metabolites with high mean abundance, 70% of which were unannotated, showed distinct profiles between mother and offspring salivary metabolomes (Supp. Fig. 7). Likewise, GC-TOF salivary metabolomes were also significantly different between mother and offspring, but far less variation was explained (PERMANOVA, R^2^= 12%, p = 0.0001, Supp. Fig. 6B). Some metabolites, such as lactulose, were much more abundant in infants and largely absent in mothers (Supp. Fig. 8). Maternal metabolomics profiles (both GC-TOF and LC-MS/MS lipidomics) have a strong individual signature, while infants do not (see PERMANOVAs, Supp, Table 2). The infant metabolome for saliva and urine had little variance attributed to which subject donated the sample, but GC-TOF was able to detect a significant change between the infant urine metabolome at birth versus 6 and 12 months of age (PERMANOVA, R^2^= 34%, p = 0.0007, Supp. Table 2). Moreover, from lipidomics data, the infant metabolome profile seemed to converge on mothers’ metabolomes as they aged, though more samples would be needed to confirm this finding (Supp. Fig. 9). For both saliva and urine, GC-TOF and lipidomics detected metabolites were more similar for mother-child pairs than for unrelated individuals (Fig. 6). Mantel tests to determine if inter-sample relationships were similar between chromatography methods (including both GC-TOF vs lipidomics) showed a strong correlation between saliva samples, and weaker correlations between urine and CVF (Supp. Table 3).

**Figure 6:**
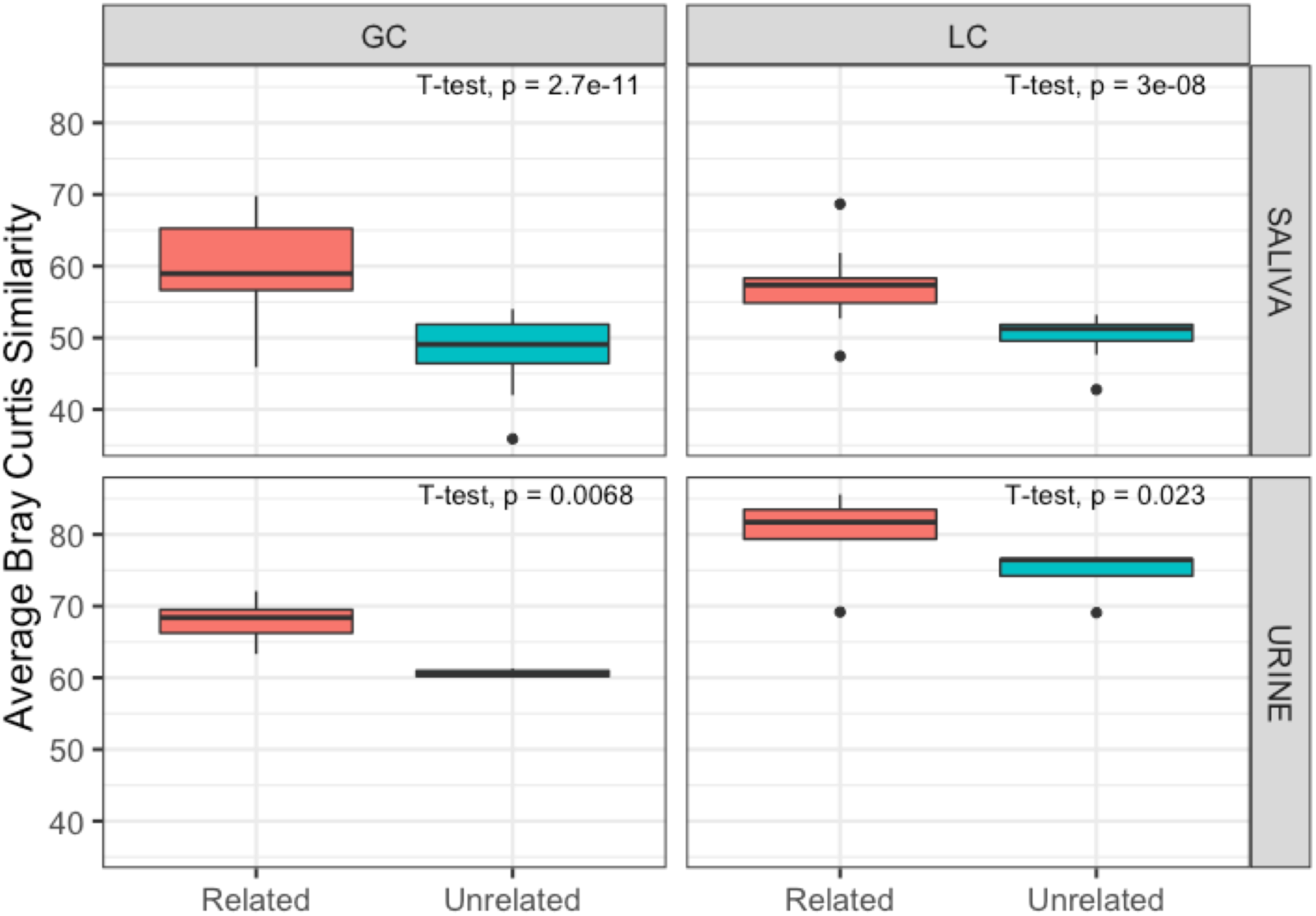
Similarity of urine and saliva metabolomes between related and unrelated individuals. Related mothers and children have significantly more similar saliva and urine metabolomes than unrelated individuals. Graph shows average bray Curtis similarity between related and unrelated individuals for GC-TOF and lipidome metabolites. Paired T-tests were done to calculate significance.

## Discussion

Exposure to the microbiome in early life is critical for immune and physiological development [1–4], yet the factors that set this trajectory remain poorly understood. In this study, we followed the vaginal microbiome through the trimesters of pregnancy for 18 women, tracking changes in the bacterial communities with longitudinal samples, and capturing their functional potential with metagenomic sequencing and multiple platforms to assess metabolomic profiles. The resolution provided by shotgun metagenomic sequencing allowed us to identify species and characterize functional gene content of CVF microbiomes. An additional strength of this work is the strict inclusion criteria defining healthy pregnancy (see *Subject Information*, in Methods). Moreover, as part of an existing sample cohort, we had the opportunity to measure saliva and urine metabolomes from mothers and children. We aim to establish how the metabolome develops in the first year of life, and how maternal-infant saliva and urine metabolomes relate. In our study, most healthy pregnant women exhibited a relatively stable vaginal microbiota throughout the trimesters of pregnancy, dominated by *Lactobacillus* or, in some cases, more diverse, *Bifidobacterium*-dominated microbiota. However, a subset of women exhibited compositional shifts in their CVF microbiota as pregnancy progressed, as has also been seen in other larger cohorts [13]. We found several strong correlations between particular vaginal communities and metabolites, which may help us understand the physiology underlying distinct vaginal microbiota structures that were evident in our study. Lastly, vaginal microbiota composition predicted which metabolites were present in the CVF samples, but not urine or saliva samples from the mothers or infants, suggesting that local microbial metabolism may represent the dominant contributor to the metabolic milieu of the vaginal tract during pregnancy.

Our study supports the results from several other studies that have indicated that the vaginal microbiome is stable during pregnancy [8, 53]. Specifically, in a longitudinal study including 90 women, most retain a microbial community with the same dominant member (in *L. crispatus* communities, 75% remain stable, in communities with high *L. iners* abundance, 71% remain stable, and in more diverse communities like those sometimes associated with BV, 58% do not shift) [13]. Using metagenomic sequencing to probe microbial community variation, our findings indicate that few bacteria, particularly *Lactobacillus* species, are highly abundant in the vaginal environment. Indeed, for individuals with vaginal microbiomes numerically dominated by *L. crispatus* or *L. iners*, the vast majority of reads mapped to contigs from one strain of *L. crispatus* or *L. iners* (Supp. Fig. 1). Of note, the microbiome of some individuals did differ considerably with advancing pregnancy. For instance, the vaginal microbiota of subjects 1180 and 1222 had higher abundances of *L. crispatus* during the first trimester, but *L. iners* was more abundant in the remaining trimesters. Brooks et al. [54] demonstrated that shifts in vaginal microbiota structures can be described probabilistically, where shifts from *L. crispatus* to *L. iners* are the most likely to occur. This is consistent with the observations made in two individuals from our study, however, due to the small number of samples exhibiting this phenomenon qualitative assessments were more appropriate than statistical analysis. Of note, vaginal microbiota instability throughout pregnancy was associated with the presence of *Candida*, a known opportunistic pathogen of the vaginal tract. Since inclusion criteria for this study stipulated no antibiotic treatment, it is unlikely that *Candida* detection was a result of antibiotic administration. The prevalence of *Candida* in our cohort is more likely to be reflective of the fact that pregnancy is a known risk factor for candidiasis [55] and to related differences in the vaginal environment, including microbiological colonization. Indeed *L. crispatus* has been shown to have anti-*Candida* activity [56], and 90% (10/11) of samples that were *Candida* positive came from individuals whose vaginal microbiota were dominated by an organism other than *L. crispatus*.

A few metabolites were highly indicative of the bacterial community present in each subject and may be useful biomarkers for the type of vaginal microbiota present. The most indicative metabolite was indole-3-lactate (ILA), a tryptophan metabolite whose abundance was correlated with communities having abundant *L. crispatus* (Fig. 4B and Supp. Fig. 4). One potential explanation is that *L. crispatus* produces ILA to competitively exclude the growth of other species (Fig. 7). At physiologically relevant concentrations, ILA has been shown to have anti-microbial properties against both Gram-positive and Gram-negative organisms [57, 58]. Although the production of lactic acid is generally thought of as a strategy *Lactobacillus* spp. use to prevent other species from colonizing the vagina, perhaps these organisms also use ILA in a similar or supplementary capacity. Additionally, bacterial derived ILA (also referred to as indole-lactic acid) has been recently shown to directly move from maternal to infant tissue [6]. It has been suggested that indoles may play an important role as a ligand for the human aryl hydrocarbon receptor (AhR), which have diverse functions from immune regulation to metabolism (reviewed in [59]). Zelante et. al further showed that some lactobacilli produce the related tryptophan catabolite, indole-3-aldehyde (IAld), which provides protection against candidiasis by increasing IL-22 production via AhR receptor binding [60]. The study also demonstrated that vaginal specific bacteria, such as *L. acidophilus*, produce IAld in the vaginal environment, which protected against vaginal but not intestinal candidiasis. We measured indole-3-acetate (IAA), the direct precursor to IAld, in our study, but found no difference in its abundance between women dominated by different species of *Lactobacillus* (data not shown). Because indole-3-lactate can act as a ligand for AhR, we hypothesize that *L. crispatus* may effectively regulate the IL-22-AhR response in the vagina, reducing the risk of vaginal candidiasis in the same way IAld does, and potentially activating the AhR response in newborns to prevent early life candidiasis.

**Figure 7:**
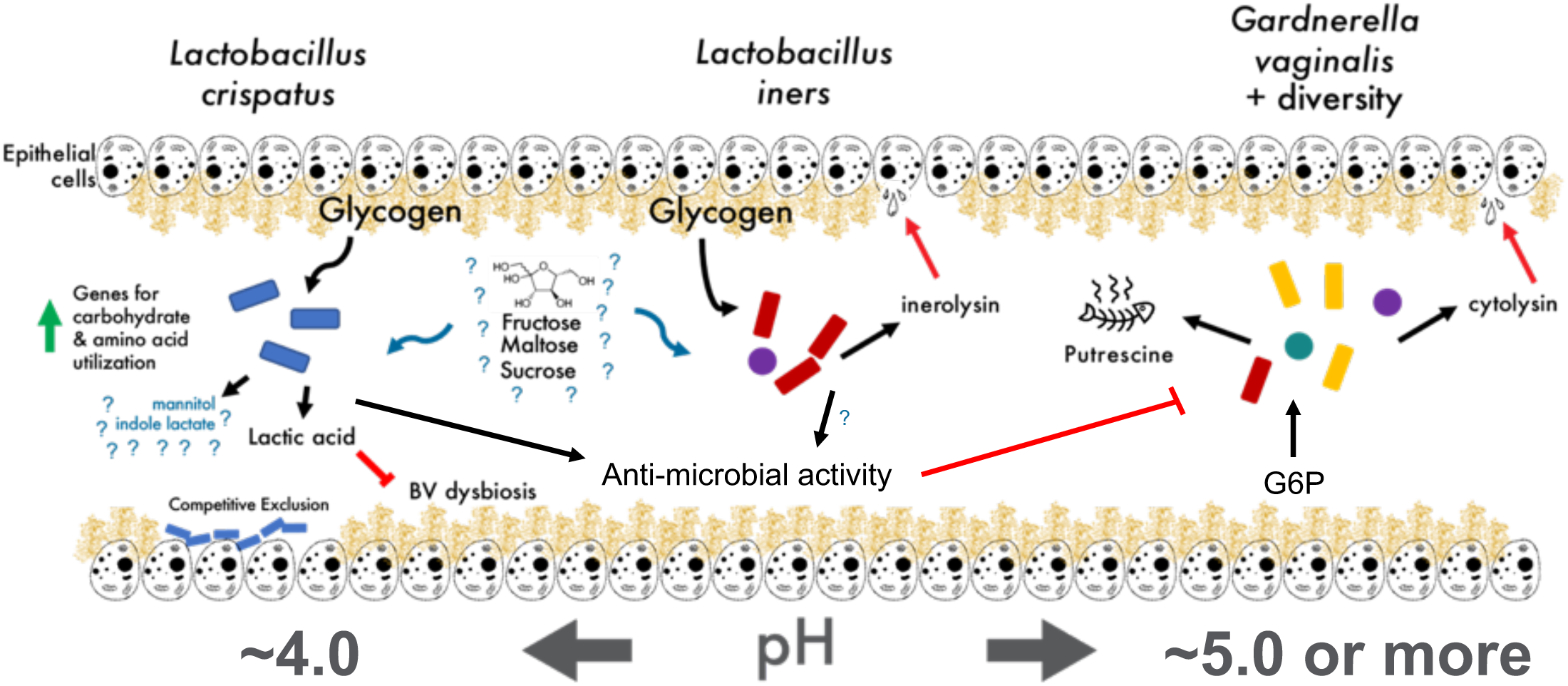
Proposed vaginal microbial community model. Current hypothesized model of vaginal microbial community physiology, with gaps in understanding (denoted by question marks) where future work is needed. Our study indicates that mannitol production is associated with a high relative abundance of *L. crispatus*.

Whole genome shotgun metagenomics allowed us to begin to address the functional capacities of these microbiomes. The largest differences between functional capacity appeared to be between communities where *L. crispatus* or *G. vaginalis* were the most abundant bacterial taxon. One pathway particularly enriched in *Gardnerella* communities was the ammonia assimilation cycle (Supp. Fig. 4). Studies have pointed out *Gardnerella’s* preference for ammonia as a nitrogen source [61]; moreover, this ability to assimilate ammonia and produce amino acids has been implicated in mutualistic interactions between species of *Prevotella*, in the context of BV [62]. Together, these BV-associated organisms contribute to genital inflammation, which may play a role in the susceptibility of certain diseases, such as HIV [63].

Increased abundance of mannitol when *L. crispatus* was present is an important and unexpected finding (Fig. 4). Mannitol itself may optimize the tonicity of the vaginal environment, and has recently been considered for this use in developing effective therapeutics for altering the vaginal microbiota [64]. Even more, mannitol may assist *L. crispatus* in adhering to the epithelial layer, a strategy the organism uses to competitively inhibit other microbes from colonizing, by drawing out excess water in the mucin layer and altering the mucin structure [65]. Irrespective of the biochemistry, these genes, and mannitol in general, represent very specific markers of a community where *L. crispatus* was most abundant.

Although it is known that homofermentative lactic acid bacteria (LAB) such as *L. crispatus* [66] convert glucose primarily to lactic acid, it is unclear why mannitol accumulates in this niche. We did find the gene mannitol-1-phosphate dehydrogenase, responsible for catalyzing the conversion of fructose to mannitol, was highly correlated (R^2^ = 0.9) with the relative abundance of *L. crispatus*. Indeed, some studies have begun to call *L. crispatus’* homofermentative classification into question [67]. Rare cases of mannitol production from homofermentative LAB usually occurred when lactate dehydrogenase was defective or knocked out [68]. Fructophilic LAB (FLAB) have been described before [69], some of which are highly efficient at producing mannitol by using fructose as both a main substrate for fermentation and an electron acceptor, yet FLAB have not been discussed in the context of the vaginal microbiome. Metabolomic analysis of our CVF samples failed to capture significant levels of fructose; however, previous studies have indicated an appreciable amount of fructose within the cervical mucus of humans [70] and the capability of *L. crispatus* to utilize fructose as a carbon source [67, 71]. Interestingly, there was no difference in the glucose abundance between the four distinct vaginal communities. One study indicated that FLAB need oxygen or pyruvate to efficiently dissimilate glucose [72]. We suggest that *L. crispatus* could represent a fructophilic LAB, which in the low abundance of oxygen and pyruvate, switches to metabolizing fructose to mannitol. This may imply that available electron acceptors play an important role in the maintenance of lactic acid in the vaginal environment, and the depletion of electron acceptors may contribute to community dysbiosis due to a rise in pH (Fig. 7). Future experiments using culturing to elucidate whether these *in vivo* community data are recapitulated with axenic cultures *in vitro* are needed.

Furthermore, this study began to investigate the utility of two metabolomic methods (GC-TOF and LC-MS/MS lipidomics) for analysis of the pregnancy and early life metabolomes. Our data showed that metabolite intensities obtained by GC-TOF were more tightly correlated with microbial community composition than those obtained by lipidomics, perhaps indicating that GC-TOF is more effective at detecting microbial metabolites than lipidomics, especially during pregnancy. We also show that both the saliva and urine metabolomes of children are more similar to their own mother than to unrelated individuals (Fig. 6). Strikingly, the ability of lipidomics to differentiate mothers from children via saliva was the strongest signal in our metabolomics data (Supp. Table 1). The oral microbiome may play a role in this, as there is a well-established community succession in children during early life (reviewed in [73]), where children begin life with oral microbes that differ from those in adults. Lactulose, detected by GC-TOF, was a specific metabolite with increased abundance in infant saliva, which may reflect its use as a treatment for constipation [74] or perhaps even its presence in heated milk [75]. Finally, urine metabolomes had a distinct age profile, especially with the lipidomics data. Our data suggests that, over the first year of life, the urine metabolome rapidly converges on the adult metabolome. This is likely a result of the development of renal system in children [76], along with the development of the gut microbiome and the related metabolites which are processed through the liver and kidneys. Other reasons for this age-related shift include a change in diet and a weaning off of breastmilk or formula [76]. Expectedly, we did not see a strong influence of the vaginal microbiome during pregnancy on the infant saliva and urine metabolome. We suspect that if differences in the vaginal microbiome were to affect the early life saliva and urine metabolome, those effects would be subtle. The lack of stool samples from the mothers and infants is a limitation of this study, as they may contain a stronger signal of shared metabolomes across mother-infant dyads. Additionally, this study explored an approach to characterize the metabolomes in high-throughput. By using acoustic deposition, in combination with MALDI-MSI, a throughput of ∼1 seconds per sample was reached using only 2 microliter of sample. Of the metabolites identified by GC-MS and LC-MS, ∼55% had corresponding ions in the MALDI-MSI analysis (Supp. Table 1). Future work will focus on confirming these metabolite identifications, but the initial results are promising and indicate that rapid analysis of microbial metabolites using MALDI, an analysis platform routinely used in clinical microbiology laboratories, is feasible [77].

Overall, we provide a broad look at the metabolome during pregnancy and early life, detailing the utility of GC-TOF, lipidomics and MALDI-MSI for saliva, urine, and CVF.

## Conclusion

Here we share a high-resolution characterization of the vaginal microbiome, longitudinally sampled throughout healthy pregnancy. We show that, despite the generally accepted view that lactobacilli are indicative of healthy vaginal communities, many women in our healthy pregnancy cohort had non-*Lactobacillus* dominated communities. The vaginal communities were characterized by a high abundance of one or a few acid-tolerant species, which dictated the physiologic potential and the metabolic profiles of the vaginal microbiome. Many of the metabolites that were specific to these different organisms warrant further investigation, especially considering the recent development of VMT as a treatment for BV [28]. The metabolites we found to be associated with *L. crispatus* may be useful as microbiome cultivation approaches are developed to intentionally direct the composition of the vaginal microbiome. For example, indole-3-lactate may support *L. crispatus* colonization, while mannitol may indicate a shift in metabolism away from fermentation and the production of acid, relaxing the low-pH selection pressure which normally gives *L. crispatus* an advantage.

## Acknowledgements

The pilot grant awarded to start a UC Center for Pediatric Microbiome Research awarded by the Institute for Clinical and Translational Science gave rise to this project. We would like to thank Prof. Dan Cooper for enthusiasm and strategic advice. We would like to thank the members of the Whiteson Lab, especially Dr. Whitney England for insight into metagenomics analysis, Joann Phan and Tara Gallagher, for thoughtful comments during the preparation of this manuscript. Dr. Heather Maughan made insightful and clarifying edits that we are grateful for.

## Data Availability

Sequence data for 16S, ITS2, and shotgun metagenomes were deposited on the National Center for Biotechnology Information (NCBI) sequence read archive (SRA) under the accession number XXXXXXXX. Metabolomics data for all samples can be found in Supplementary Table 1. R-scripts for statistical analysis are published on GitHub: https://github.com/aoliver44/Cervicovaginal-Paper (embargoed until publication).

## Ethics approval and consent to participate

This study utilized a subset of samples from a larger, longitudinal prospective cohort study designed to analyze the relationship between maternal stress and infant development, conducted at the University of California, Irvine (UCI) [30]. The University of California’s Institutional Review Board approved the protocol and written, informed consent was obtained from each participant. Research on human subjects was performed in accordance with the Declaration of Helsinki.

## Consent for publication

Not applicable.

## Figure Legends

Supp. Fig. 1: Anvi’o plot illustrating the abundance of short (>= 2,000bp) contigs for each subject (trimesters combined). The participants are colored according to the most abundant microbe present (*L. iners* (blue), *L. crispatus* (green), *G. vaginalis* (yellow)). Short read taxonomic assigner Kaiju was used to assign taxonomy to the contigs

Supp. Fig. 2: A) Bar plot showing the relative abundance of fungal OTUs within subjects. The most abundant OTUs by far were all classified as *Candida albicans* (pink). All other OTUs made up very little of the fungal presence in these samples, save for subject 1088, who had a species of *Aspergillus* as its most abundant fungal taxon. B) Sample where *Candida* was detected had greater shifts in bacterial composition than samples without.

Supp. Table 1: A spreadsheet of the raw metabolic data, separated by tabs. Metadata is also included.

Supp. Fig. 3: Lefse analysis of the HUMAnN2 output, specifically the mannitol degradation pathway. Pairwise t-tests of *L. crispatus* against other dominant microbes.

Supp. Fig. 4: A permutated random forest recapitulates the metabolites that drive differences between the vaginal communities. The heatmap shows the mean decrease in accuracy associated with the specific microbiomes. Bold boxes around the heatmap cell indicates statistical significance of that feature at p < 0.05.

Supp. Fig. 5: Lefse analysis of annotated HUMAnN2 output comparing the enrichment of functional pathways between vaginal microbiomes with abundant *Bifidobacteriaceae (G. vaginalis), L. crispatus, L. iners*, or other. This analysis was done using the -no_stratify flag for humann2, which analyzes the data without the taxa specific information.

Supp. Table 2: PERMANOVA, from the Primer-e software, analysis of participant, dyad, timepoint, and most abundant microbe on the metabolomes of saliva, urine, and cervical vaginal fluid. Results are separated based on chromatography was used. In order to calculate p-values, 9999 permutations were used. Both p-values and monte-carlo p-values are presented. ECoV were calculated by dividing the component of variation by the sum total of all components of variation.

Supp. Table 3: RELATE test, from the Primer-e software, which calculates how shared among-sample relationships are between two distance matrices (analogues to Mantel coefficients). Higher rho values, and significance values, indicate the relationship between samples are similar in both distance matrices. All samples from mother and child were used in the construction of the Bray Curtis dissimilarity matrices.

Supp. Fig. 6: A) Principle coordinates ordination of saliva LC-MS/MS lipidomics, colored by whether the sample originated from mother or child. B) Principle coordinates ordination of saliva metabolomes by GC-TOF, colored by whether the sample originated from mother or child.

Supp. Fig. 7: Analysis of the top 50 lipidomic metabolites from saliva that had the highest mean abundance across all samples using Primer-e software. Metabolites were initially standardized within each sample, and then across variables. Color indicates this normalized abundance of the metabolite. Where present, annotations were used for lipidomic data.

Supp. Fig. 8: Analysis of the top 50 lipidomic metabolites from saliva that had the highest mean abundance across all samples using Primer-e software. Metabolites were initially standardized within each sample, and then across variables. Color indicates this normalized abundance of the metabolite. When present, annotations were used for GC-TOF data, otherwise BinBase numbers are shown.

Supp. Fig. 9: Principle coordinates ordination of urine lipidomes, colored by age. All timepoints for mothers were used (i.e. Trimesters 1-3).

